# Reference genome of the bicolored carpenter ant, *Camponotus vicinus*

**DOI:** 10.1101/2023.06.26.546281

**Authors:** Philip S. Ward, Elizabeth I. Cash, Kailey Ferger, Merly Escalona, Ruta Sahasrabudhe, Courtney Miller, Erin Toffelmier, Colin Fairbairn, William Seligmann, H. Bradley Shaffer, Neil D. Tsutsui

## Abstract

Carpenter ants in the genus Camponotus are large, conspicuous ants that are abundant and ecologically influential in many terrestrial ecosystems. The bicolored carpenter ant, C. vicinus Mayr, is distributed across a wide range of elevations and latitudes in western North America, where it is a prominent scavenger and predator. Here, we present a high-quality genome assembly of C. vicinus from a sample collected in Sonoma County, CA, near the type locality of the species. This genome assembly consists of 38 scaffolds spanning 302.74 Mb, with contig N50 of 15.9Mb, scaffold N50 of 19.9 Mb, and BUSCO completeness of 99.2%. This genome sequence will be a valuable resource for exploring the evolutionary ecology of C. vicinus and carpenter ants generally. It also provides an important tool for clarifying cryptic diversity within the C. vicinus species complex, a genetically diverse set of populations, some of which are quite localized and of conservation interest.

## Introduction

The ant tribe Camponotini contains almost 2000 described species, of which a little more than half belong to *Camponotus*, the world’s most widely distributed ant genus (Bolton 2023). Many species of *Camponotus* nest in rotting wood, earning them the common name “carpenter ants” (Hansen & Klotz 2005). All species of Camponotini harbor obligate, vertically-inherited gut bacteria (*Blochmannia*) that provide important nutritional benefits and likely contribute to host survival under varying environmental conditions (Feldhaar *et al*. 2007; Williams & Wernegreen 2015). Some *Camponotus* ants are also common structural pests, causing costly damage as they excavate wooden structures.

Carpenter ants in the *Camponotus vicinus* species complex are prominent scavenging and predatory ants, occurring in all ecoregions of California except the Colorado and Sonoran Deserts. In higher elevation conifer forests of California, *C. vicinus* commonly nests in and around fallen, decomposing logs, and is one of the most abundant ground-dwelling arthropods (Figure 1A). This complex includes two widespread species as well as several cryptic taxa with more limited distributions that are of conservation interest. The cryptic diversity in the *C. vicinus* complex includes an undescribed species endemic to the Channel Islands.

**Figure 1.**
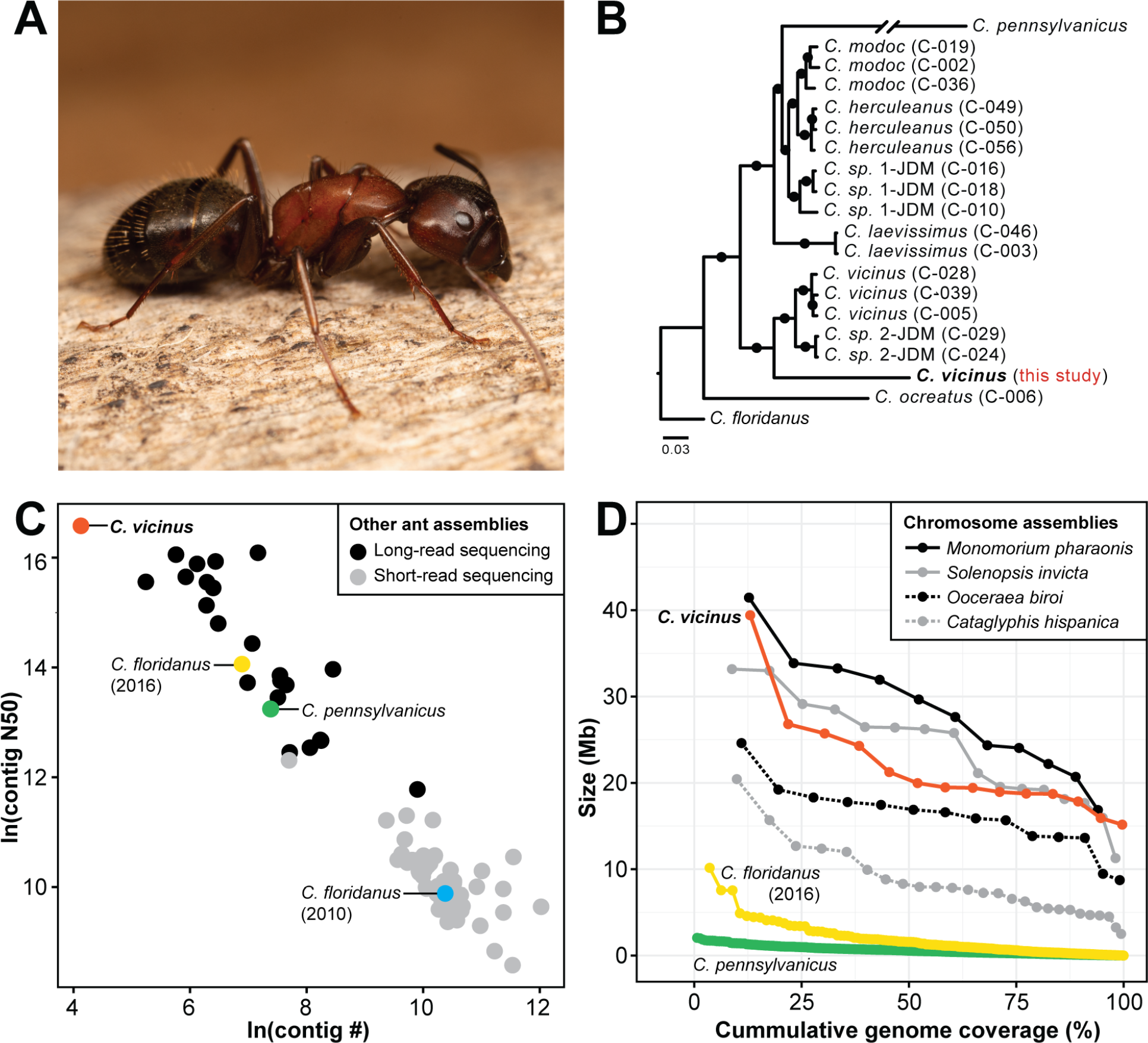
Bicolored carpenter ant reference genome assembly. (**A**) A major worker of the bicolored carpenter ant, *Camponotus vicinus* (photo: Elizabeth Cash). (**B**) Phylogenetic reconstruction based on whole genome sequences of *C. vicinus* (California, this study) compared with nine other *Camponotus* species from Shields *et al*. (2018), Manthey *et al*. (2022), and Faulk (2023). Filled circles represent 100% bootstrap support. Sample names from Manthey *et al*. (2022) are shown in parentheses. (**C**) Scatterplot comparing *C. vicinus* genome assembly (red) to assemblies of *C. floridanus* (yellow = 2016 assembly, Shield *et al*. 2018; blue = 2010 assembly Bonasio *et al*. 2010), *C. pennsylvanicus* (green, Faulk 2023), and other non-*Camponotus* ant species (black = long-read sequencing, grey = short-read sequencing; n=81 total assemblies representing 59 species) based on the natural log (ln) of contig number and contig N50 values. (**D**) Lineplot comparing scaffold/contig sizes (Mb) and cumulative genome coverage (%) for *C. vicinus* (red, scaffolds), *C. floridanus* (2016, yellow, scaffolds), and *C. pennsylvanicus* (green, contigs) genome assemblies along with 4 representative ant genomes with chromosome-level assemblies (*Cataglyphis hispanica* [grey, dashed], *Monomorium pharonsis* [black, solid], *Ooceraea biroi* [black, dashed], and *Solenopsis invicta* [grey, solid]).

We report here a high-quality *de novo* reference genome assembly for *C. vicinus* collected near the type locality for this species. Existing genomic resources include an annotated reference genome for the relatively distantly related *C. floridanus* (Bonasio *et al*. 2010, Shields *et al*. 2018), as well as more recent genome sequences from *C. pennsylvanicus* (Faulk 2023) and several species collected in the American Southwest (including putative *C. vicinus* from Arizona) (Manthey *et al*. 2022). We also reconstruct a phylogeny using these *C. vicinus* genomes and several other *Camponotus* species from Manthey *et al*. (2022).

## Methods

### Biological Materials

A large, populous colony of *Camponotus vicinus*, containing a single dealate queen, numerous workers, alate queens, alate males, eggs, larvae, and pupae, was located near the type locality of this species. Collection data are as follows: USA, California, Sonoma County, 6 km east of Mark West Springs, 365 m elevation, 38.54192°N 122.64803°W, 24 July 2021, ex rotten log in *Pseudotsuga*-*Quercus* forest, P. S. Ward collector, collection code PSW18465. A worker voucher specimen from this colony, assigned the unique specimen code CASENT0886928, has been deposited in the Bohart Museum of Entomology, University of California, Davis. Workers from the sampled colony agree closely in color, pilosity, and pubescence with a syntype worker of *C. vicinus* from Calistoga, California, illustrated on AntWeb (www.antweb.org), under specimen code CASENT0915806. Our collection site is 7 km southwest of Calistoga. From the sampled colony, a single male pupa was used for HiFi sequencing and a single adult male was used for the Omni-C library.

### High molecular weight DNA extraction and nucleic acid library preparation

Flash frozen male pupa was homogenized in 650µl of homogenization buffer (10mM Tris-HCL-pH 8.0 and 25mM EDTA) using TissueRuptor II (Qiagen, Germany; Cat # 9002755). 650µl of lysis buffer (10mM Tris, 25mM EDTA, 200mM NaCl, and 1% SDS) and proteinase K (100µg/ml) were added to the homogenate and it was incubated overnight at room temperature. Lysate was treated with RNAse A (20µg/ml) at 37 ºC for 30 minutes and was cleaned with equal volumes of phenol/chloroform using phase-lock gels (Quantabio, Beverly, MA; Cat # 2302830). The DNA was precipitated by adding 0.4X volume of 5M ammonium acetate and 3X volume of ice-cold ethanol. The DNA pellet was washed twice with 70% ethanol and resuspended in an elution buffer (10mM Tris, pH 8.0). DNA was further cleaned with Zymo gDNA clean and concentrator kit (Zymo Research, Irvine, CA; Cat # 4033). To retain large DNA fragments, columns from large fragment DNA recovery kit (Zymo Research, Cat # D4045) were used during purification. Purity of gDNA was accessed using NanoDrop ND-1000 spectrophotometer where 260/280 ratio of 1.8 and 260/230 ratio of 2.26 was observed. DNA was quantified by Qubit 2.0 Fluorometer (Thermo Fisher Scientific, Waltham, MA) and total yield of 1.5µg was obtained. Integrity of the HMW gDNA was verified on a Femto pulse system (Agilent Technologies, Santa Clara, CA) where 73% of DNA was observed in fragments above 50 Kb.

The HiFi SMRTbell library was constructed using the SMRTbell Express Template Prep Kit v2.0 (Pacific Biosciences -PacBio, Menlo Park, CA, Cat. #100-938-900) according to the manufacturer’s instructions. HMW gDNA was sheared to a target DNA size distribution between 12-20 kb. The sheared gDNA was concentrated using 1.8X of AMPure PB beads (PacBio,Cat. #100-265-900) for the removal of single-strand overhangs at 37°C for 15 minutes, followed by further enzymatic steps of DNA damage repair at 37°C for 30 minutes, end repair and A-tailing at 20°C for 10 minutes and 65°C for 30 minutes, and ligation of overhang adapter v3 at 20°C for 60 minutes. The SMRTbell library was purified and concentrated with 0.45X Ampure PB beads for size selection with 40% diluted AMPure PB beads (PacBio, Cat. #100-265-900) to remove short SMRTbell templates <3 kb. The 12-20 kb average HiFi SMRTbell library was sequenced at UC Davis DNA Technologies Core (Davis, CA) using two 8M SMRT cells, Sequel II sequencing chemistry 2.0, and 30-hour movies each on a PacBio Sequel II sequencer.

The Omni-C library was prepared using the Dovetail™ Omni-C™ Kit (Dovetail Genomics, Scotts Valley, CA) according to the manufacturer’s protocol with slight modifications. First, specimen tissue (whole adult male, ID: PSW18465-M) was thoroughly ground with a mortar and pestle while cooled with liquid nitrogen. Subsequently, chromatin was fixed in place in the nucleus. The suspended chromatin solution was then passed through 100 µm and 40 µm cell strainers to remove large debris. Fixed chromatin was digested under various conditions of DNase I until a suitable fragment length distribution of DNA molecules was obtained. Chromatin ends were repaired and ligated to a biotinylated bridge adapter followed by proximity ligation of adapter containing ends. After proximity ligation, crosslinks were reversed, and the DNA was purified from proteins. Purified DNA was treated to remove biotin that was not internal to ligated fragments. An NGS library was generated using an NEB Ultra II DNA Library Prep kit (NEB, Ipswich, MA) with an Illumina compatible y-adaptor. Biotin-containing fragments were then captured using streptavidin beads. The post-capture product was split into two replicates prior to PCR enrichment to preserve library complexity with each replicate receiving unique dual indices. The library was sequenced at Vincent J. Coates Genomics Sequencing Lab (Berkeley, CA) on an Illumina NovaSeq 6000 platform (Illumina, San Diego, CA) to generate approximately 100 million 2 × 150 bp read pairs per GB genome size.

### DNA Sequencing and Genome Assembly

#### Nuclear genome Assembly

We assembled the genome of *C. vicinus* following the CCGP assembly pipeline Version 5.1, as outlined in Table 1, which lists the tools and non-default parameters used in the assembly. The pipeline uses PacBio HiFi reads and Omni-C data to produce high quality and highly contiguous genome assemblies. First, we removed the remnant adapter sequences from the PacBio HiFi dataset using HiFiAdapterFilt (Sim *et al*. 2022) and generated the initial haploid assembly using HiFiasm (Cheng *et al*. 2021) with the filtered PacBio HiFi reads and the Omni-C dataset. This process generated multiple assemblies and we kept the output assembly tagged as haplotype 1 given the ploidy of the species. We then aligned the Omni-C data to the assembly following the Arima Genomics Mapping Pipeline (https://github.com/ArimaGenomics/mapping_pipeline) and then scaffolded it with SALSA (Ghurye *et al*. 2017, Ghurye *et al*. 2019).

**Table 1:** Assembly Pipeline and Software Used.

| Table 1: Assembly Pipeline and Software Used. |  |  |
| --- | --- | --- |
| Nuclear assembly | Software and options | Version |
| Filtering PacBio HiFi adapters | HiFiAdapterFilt | Commit 64d1c7b |
| K-mer counting | Meryl (k=21) | 1 |
| Estimation of genome size and heterozygosity | GenomeScope | 2 |
| *De novo* assembly (contiging) | HiFiasm (Hi-C Mode, -primary, output p_ctg.hap1, p_ctg.hap2) | 0.16.1-r375 |
| Scaffolding |  |  |
| Omni-C data alignment | Arima Genomics Mapping Pipeline | Commit 2e74ea4 |
| Omni-C Scaffolding | SALSA (-DNASE, -i 20, -p yes) | 2 |
| Gap closing | YAGCloser (-mins 2 -f 20 -mcc 2 -prt 0.25 -eft 0.2 -pld 0.2) | Commit 0e34c3b |
| Omni-C Contact map generation |  |  |
| Short-read alignment | BWA-MEM (-5SP) | 0.7.17-r1188 |
| SAM/BAM processing | samtools | 1.11 |
| SAM/BAM filtering | pairtools | 0.3.0 |
| Pairs indexing | pairix | 0.3.7 |
| Matrix generation | cooler | 0.8.10 |
| Matrix balancing | hicExplorer (hicCorrectmatrix correct --filterThreshold -2 4) | 3.6 |
| Contact map visualization | HiGlass | 2.1.11 |
|  | PretextMap | 0.1.4 |
|  | PretextView | 0.1.5 |
|  | PretextSnapshot | 0.0.3 |
| **Genome quality assessment** |  |  |
| Basic assembly metrics | QUAST (--est-ref-size) | 5.0.2 |
| Assembly completeness | BUSCO (-m geno, -l insecta) | 5.0.0 |
|  | Mercury | 2020-01-29 |
| **Contamination screening** |  |  |
| Local alignment tool | BLAST+ (-db nt, -outfmt '6 qseqid staxids bitscore std' , -max_target_seqs 1, -max_hsps 1, -evalue 1e-25 ) | 2.1 |
| General contamination screening | BlobToolKit | 2.3.3 |
| **Endosymbiont assembly** |  |  |
| Sequence alignment | lastz (--nogapped --notransition --step=20 --format=lav) | 1.04.15 |
| Sequence alignment visualization | laj | 2005-12-14 |
| Long-read alignment | minimap2 (-ax map-pb) | 2.24-r1122 |
| SAM/BAM processing | samtools (view -hSb -F4 -F0x800) | 1.11 |
| de novo assembly of endosymbiont | HiFiasm (--primary) | 0.16.1-r375 |
| Extraction of PacBio HiFi reads from alignment | seqtk (subseq) | 1.3-r117-dirty |
| Annotation of the bacterial genome | bakta | 1.7.0 (DB: 5.0.0) |
| **Assembly comparisons** |  |  |
| Data visualization | R (ggplot2) | v4.3.0 (v3.3.6) |
| **Species** | **Accession #** | **Reference** |
| *Camponotus floridanus* | GCA_003227725.1 | Shields et al. 2018 |
| *Camponotus pennsylvanicus* | GCA_023638675.1 | Faulk 2023 |
| *Cataglyphis hispanica* | GCA_021464435.1 | Darras et al. 2022 |
| *Monomorium pharaonis* | GCA_013373865.2 | Gao et al. 2020 |
| *Ooceraea biroii* | GCA_003672135.1 | McKenzie and Kronauer 2018 |
| *Solenopsis invicta* | GCA_016802725.1 | Helleu et al. 2022 |
| **Phylogenetic analysis** |  |  |
| Quality filtering, adapter trimming | bbmap (bbduk.sh) | v39.01 |
| Alignment to reference | BWA (BWA-MEM) | v0.7.17 |
| Sort files, identify duplicates | PicardTools (SortSam, MarkDuplicates) | v1.141 |
| Alignment metrics, read depth | samtools (flagstat, depth, index) | v1.8 |
| Genome alignment | MUMmer (nucmer, --sam-long) | v4.0.0rc1 |
| Variant calling | BCFtools (mpileup, call) | v1.6 |
| Quality filtering | VCFtools | v0.1.15 |
| Model selection | jModelTest | v2.1.10 |
| Phylogenetic reconstruction | RAxML (best tree -f a) | v8.2.12 |

The genome assembly was manually curated by iteratively generating and analyzing their corresponding Omni-C contact maps. To generate the contact maps we aligned the Omni-C data with BWA-MEM (Li 2013), identified ligation junctions, and generated Omni-C pairs using pairtools (Golobordko *et al*. 2018). We generated a multi-resolution Omni-C matrix with cooler (Abdennur an Mirny 2020) and balanced it with hicExplorer (Ramírez *et al*. 2018). We used HiGlass (Kerpedjiev *et al*. 2018) and the PretextSuite (https://github.com/wtsi-hpag/PretextView; https://github.com/wtsi-hpag/PretextMap; https://github.com/wtsi-hpag/PretextSnapshot) to visualize the contact maps where we identified misassemblies and misjoins, and finally modified the assembly using the Rapid Curation pipeline from the Wellcome Trust Sanger Institute, Genome Reference Informatics Team (https://gitlab.com/wtsi-grit/rapid-curation). Some of the remaining gaps (joins generated during scaffolding and/or curation) were closed using the PacBio HiFi reads and YAGCloser (https://github.com/merlyescalona/yagcloser). Finally, we checked for contamination using the BlobToolKit Framework (Challis *et al*. 2020).

#### Genome assembly assessment

We generated k-mer counts from the PacBio HiFi reads using meryl (https://github.com/marbl/meryl). The k-mer counts were then used in GenomeScope2.0 (Ranallo-Benavidez *et al*. 2020) to estimate genome features including genome size, heterozygosity, and repeat content. To obtain general contiguity metrics, we ran QUAST (Gurevich *et al*. 2013). To evaluate genome quality and functional completeness we used BUSCO (Manni *et al*. 2021) with the Arthropoda ortholog database (arthropoda_odb10) which contains 1,013 genes. Assessment of base level accuracy (QV) and k-mer completeness was performed using the previously generated meryl database and merqury (Rhie *et al*. 2020). We further estimated genome assembly accuracy via BUSCO gene set frameshift analysis using the pipeline described in Korlach *et al*. (2017). Measurements of the size of the phased blocks is based on the size of the contigs generated by HiFiasm. We follow the quality metric nomenclature established by Rhie *et al*. (2021), with the genome quality code x.y.P.Q.C, where, x = log10[contig NG50]; y = log10[scaffold NG50]; P = log10 [phased block NG50]; Q = Phred base accuracy QV (quality value); C = % genome represented by the first ‘n’ scaffolds, based on a karyotype of n=14, reported in the related species, *C. ligniperdus* (Hauschteck & Jungen 1983) and *C. japonicus* (Imai 1966).

#### Endosymbiont genome assembly

We used the genome of *Blochmannia* (NCBI:GCF_023585685.1; ASM2358568v1; Manthey, Giron and Hruska 2022) as a guide to assemble the endosymbiont genome present in our sample. We aligned the contigs that were removed from the nuclear genome in the contamination process to the ASM2358568v1 reference using lastz (Harris 2007) to verify existence of the endosymbiont in the assembly. We aligned the adapter-trimmed PacBio HiFi reads to the *Blochmannia* sequence using minimap2 (Li 2018; 2021) and samtools (Danecek *et al*. 2021), and filtered out secondary alignments, unmapped reads and reads that failed platform/vendor quality checks. We extracted the reads left from the alignment and used them to *de novo* assemble a *Blochmannia* genome with HiFiasm. Finally, we used bakta (Schwengers *et al*. 2021; (https://bakta.computational.bio/) to generate a draft genome annotation of the bacterial genome to assess completeness of the genome.

#### Assembly comparisons

We compared basic nuclear genome assembly metrics for all 59 ant species currently available in GenBank using NCBI assembly reports (Table S1). Contig number versus contig N50 (both ln transformed) results were plotted using ggplot2 in R (Wickham 2016; Figure 1C) to visualize differences in contiguity between ant genomes. Additionally, scaffold and chromosome sizes (Mb) were plotted relative to genome coverage (%) for four ant species with chromosome-level assemblies (*Cataglyphis hispanica, Monomorium pharaonis, Ooceraea biroi*, and *Solenopsis invicta*) along with three *Camponotus* species, *C. vicinus* (this study), *C. floridanus* (Shields *et al*. 2018), and *C. pennsylvanicus* (Faulk 2023), to compare contiging and scaffolding results among genome assemblies (Figure 1D, Table S2).

#### Phylogenetic analysis

Our dataset for phylogenetic analysis consisted of 17 whole-genome sequencing (WGS) samples described in Manthey *et al*. (2022), a *Camponotus pennsylvanicus* reference genome, our assembled *Camponotus vicinus* reference genome, and the *Camponotus floridanus* reference genome which served as our outgroup (NCBI BioProjects PRJNA839641, PRJNA820489, PRJNA874059, and PRJNA476946, respectively). We performed quality filtering and adapter trimming of the sequencing reads from the 17 WGS samples with the bbduk.sh script from the bbmap package (Bushnell 2014). We then aligned these samples to the *C. floridanus* reference genome with the BWA-MEM. We used PicardTools (Broad Institute 2019) to sort our resulting SAM files and flag duplicates using the SortSam and MarkDuplicates commands. We also computed alignment metrics and read depth, as well as built bam indexes using the samtools (Li *et al*. 2009) flagstat, depth, and index commands. The assembled reference genomes were aligned to the *C. floridanus* reference genome using the MUMmer (Marçais *et al*. 2018) alignment tool. The resulting sam files were reformatted using an in-house bash script to follow the proper input formatting for samtools. Finally, these files were first sorted by read group and then converted to BAM format using the samtools sort and samtools view -b commands. We performed variant calling with BCFtools (Li 2011) for all samples using the mpileup and call commands. We then performed quality filtering with VCFtools (Danecek *et al*. 2011), removing sites with the following specifications: minor allele frequency (MAF) <0.05, missing in >25% of samples, quality score <30, and read depth <10 or >100.

We converted our VCF file to phylip alignment format using the python script vcf2phylip.py (Ortiz 2019). We used RAxML (Stamatakis 2014) to generate our phylogenetic tree by performing a best tree search (option -f a) with 1000 rapid bootstrap replicates (option -x). We determined the ‘best-fit’ model of nucleotide substitution to be GTR using jModelTest (Darriba 2012, Guindon & Gascuel 2003).

## Results

### Sequencing data

The Omni-C and PacBio HiFi sequencing libraries generated 18.29 million read pairs and 1.4 million reads, respectively. The latter yielded 52.19 fold coverage (N50 read length 12,799 bp; minimum read length 54 bp; mean read length 11,675 bp; maximum read length of 58,419 bp) based on the Genomescope 2.0 genome size estimation of 315 Mb. Based on PacBio HiFi reads, we estimated 0.129% sequencing error rate. The k-mer spectrum based on PacBio HiFi reads show (Figure 2A) a unimodal distribution with a single peak at ∼51.

**Figure 2.**
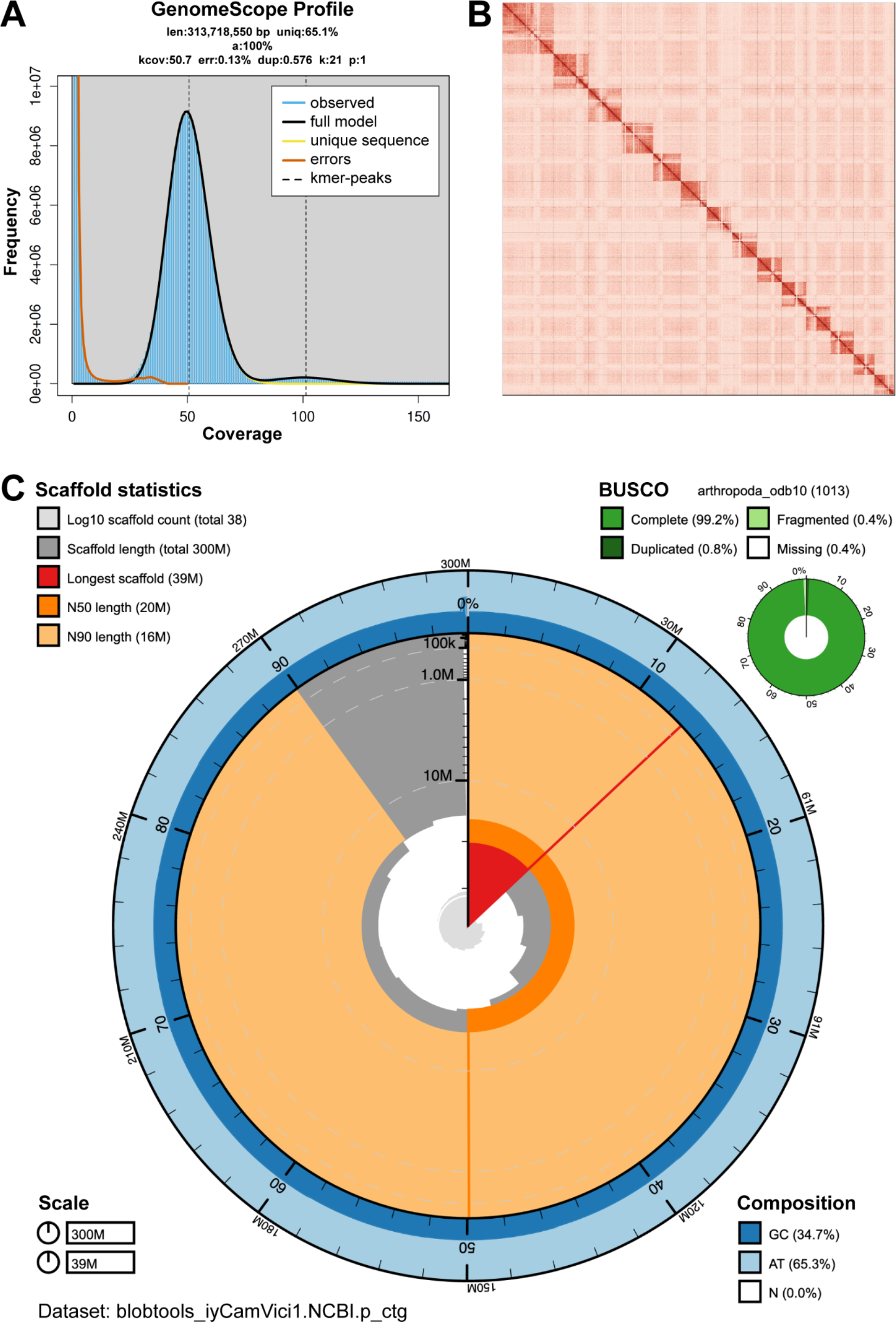
Visual overview of genome assembly metrics. (**A**) K-mer spectra output generated from PacBio HiFi data without adapters using GenomeScope2.0. The unimodal pattern observed corresponds to a haploid genome. (**B**) Omnii-C Contact map for the genome assembly generated with PretextSnapshot. The Omni-C contact map translates proximity of genomic regions in 3-D space to contiguous linear organization. Each cell in the contact map corresponds to sequencing data supporting the linkage (or join) between two of such regions. Scaffolds are separated by black lines and higher density corresponds to higher levels of fragmentation. (**C**) BlobToolKit Snail plot showing a graphical representation of the quality metrics presented in Table 2 for the *Camponotus vicinus* primary assembly. The plot circle represents the full size of the assembly. From the inside to the outside, the central plot covers length-related metrics. The red line represents the size of the longest scaffold; all other scaffolds are arranged in size order moving clockwise around the plot and drawn in gray starting from the outside of the central plot. Dark and light orange arcs show the scaffold N50 and scaffold N90 values. The central light gray spiral shows the cumulative scaffold count with a white line at each order of magnitude. White regions in this area reflect the proportion of Ns in the assembly. The dark vs. light blue area around it shows mean, maximum and minimum GC vs. AT content at 0.1% intervals.

### Nuclear genome assembly

The final assembly (iyCamVici1) genome size is close to the estimated value from Genomescope2.0 (Figure 2A, Pflug *et al*. 2020). The assembly consists of 38 scaffolds spanning 302.74 Mb with contig N50 of 15.9Mb, scaffold N50 of 19.9 Mb, longest contig of 22.35 Mb and largest scaffold of 39.41 Mb. Detailed assembly statistics are reported in tabular form in Table 2, and graphical representation for the assembly in Figure 2B. The iyCamVici1 assembly has a BUSCO completeness score of 99.2% using the Arthropoda gene set, a per-base quality (QV) of 68.45, a k-mer completeness of 99.41 and a frameshift indel QV of 54.57.

**Table 2:**
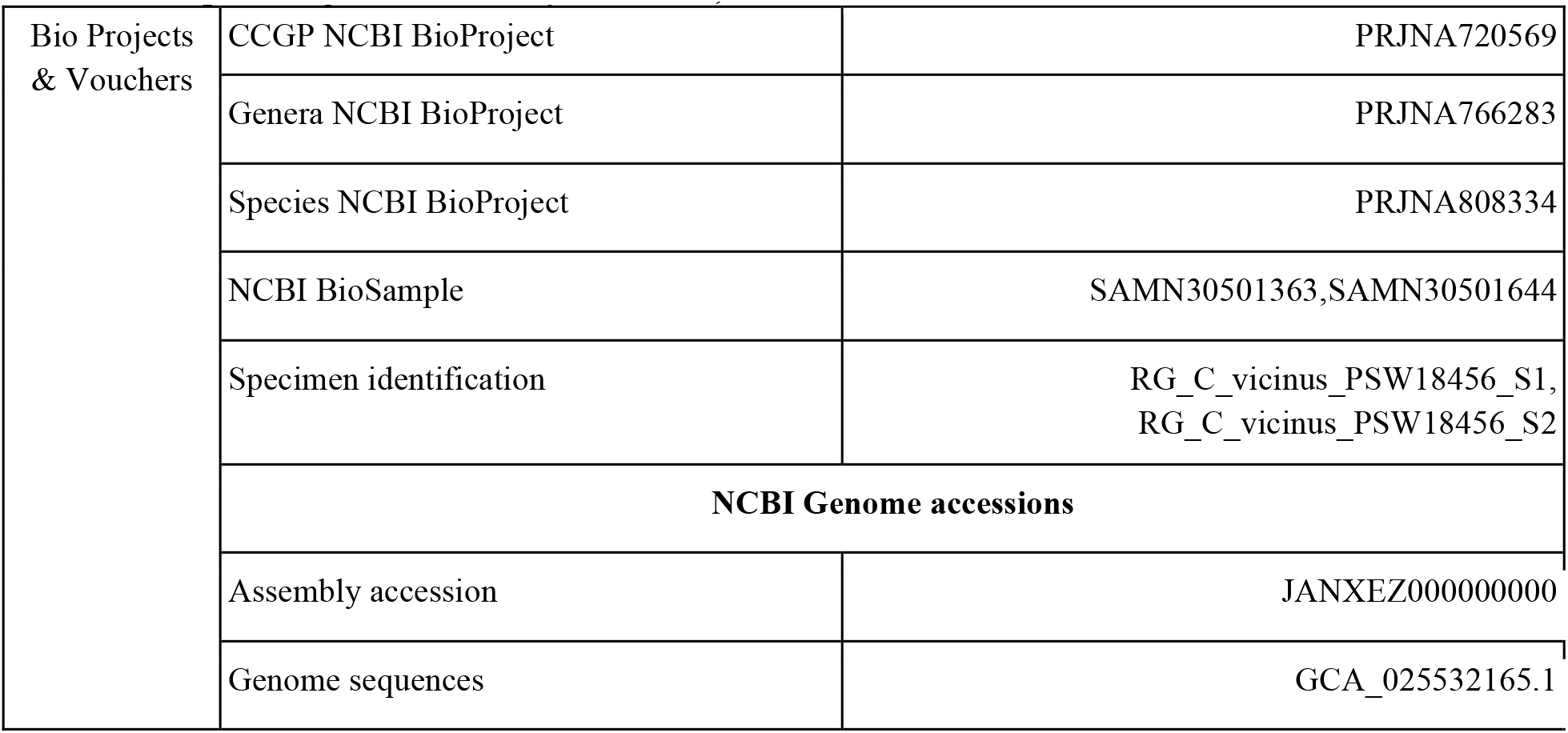
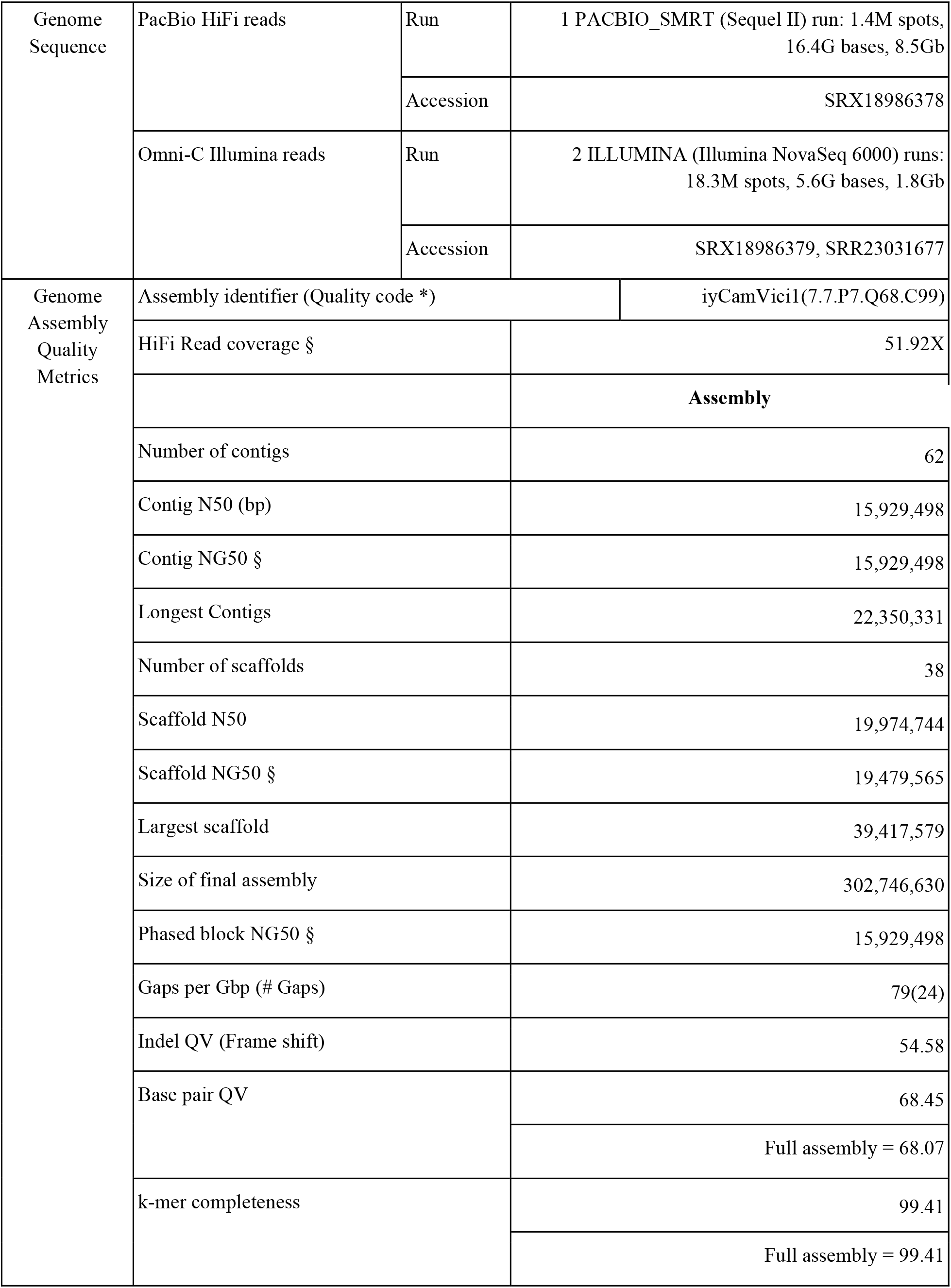
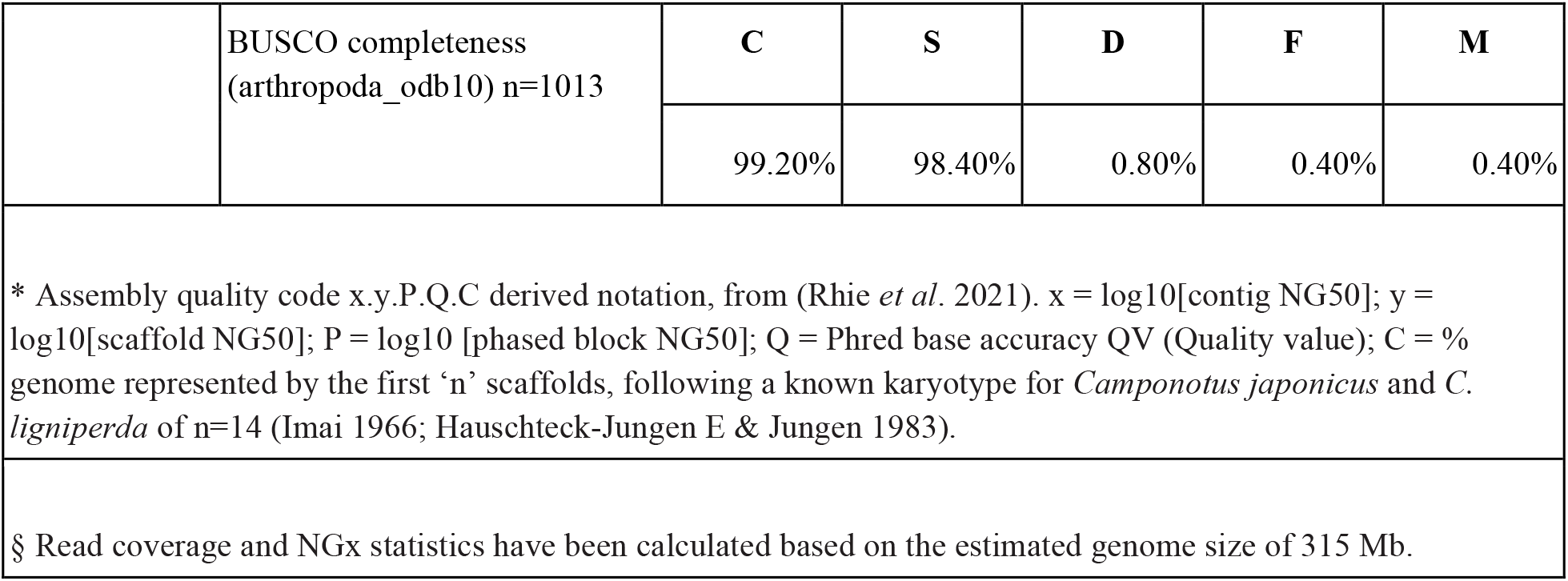
Sequencing and assembly statistics, and accession numbers.

|  |  |  |
| --- | --- | --- |
| Bio Projects & Vouchers | CCGP NCBI BioProject | PRJNA720569 |
|  | Genera NCBI BioProject | PRJNA766283 |
|  | Species NCBI BioProject | PRJNA808334 |
|  | NCBI BioSample | SAMN30501363,SAMN30501644 |
|  | Specimen identification | RG_C_vicinus_PSW18456_S1,<br>RG_C_vicinus_PSW18456_S2 |
|  | **NCBI Genome accessions** |  |
|  | Assembly accession | JANXEZ000000000 |
|  | Genome sequences | GCA_025532165.1 |

|  |  |  |  |
| --- | --- | --- | --- |
| Genome Sequence | PacBio HiFi reads | Run | 1 PACBIO_SMRT (Sequel II) run: 1.4M spots, 16.4G bases, 8.5Gb |
|  |  | Accession | SRX18986378 |
|  | Omni-C Illumina reads | Run | 2 ILLUMINA (Illumina NovaSeq 6000) runs: 18.3M spots, 5.6G bases, 1.8Gb |
|  |  | Accession | SRX18986379, SRR23031677 |
| Genome Assembly Quality Metrics | Assembly identifier (Quality code *) |  | iyCamVici1(7.7.P7.Q68.C99) |
|  | HiFi Read coverage § |  | 51.92X |
|  |  |  | **Assembly** |
|  | Number of contigs |  | 62 |
|  | Contig N50 (bp) |  | 15,929,498 |
|  | Contig NG50 § |  | 15,929,498 |
|  | Longest Contigs |  | 22,350,331 |
|  | Number of scaffolds |  | 38 |
|  | Scaffold N50 |  | 19,974,744 |
|  | Scaffold NG50 § |  | 19,479,565 |
|  | Largest scaffold |  | 39,417,579 |
|  | Size of final assembly |  | 302,746,630 |
|  | Phased block NG50 § |  | 15,929,498 |
|  | Gaps per Gbp (# Gaps) |  | 79(24) |
|  | Indel QV (Frame shift) |  | 54.58 |
|  | Base pair QV |  | 68.45 |
|  |  |  | Full assembly = 68.07 |
|  | k-mer completeness |  | 99.41 |
|  |  |  | Full assembly = 99.41 |

|  |  |  |  |  |  |  |
| --- | --- | --- | --- | --- | --- | --- |
|  | BUSCO completeness<br>(arthropoda_odb10) n=1013 | **C** | **S** | **D** | **F** | **M** |
|  |  | 99.20% | 98.40% | 0.80% | 0.40% | 0.40% |
| * Assembly quality code x.y.P.Q.C derived notation, from (Rhie *et al.* 2021). x = log10[contig NG50]; y = log10[scaffold NG50]; P = log10 [phased block NG50]; Q = Phred base accuracy QV (Quality value); C = % genome represented by the first 'n' scaffolds, following a known karyotype for *Camponotus japonicus* and *C. ligniperda* of n=14 (Imai 1966; Hauschteck-Jungen E & Jungen 1983). |  |  |  |  |  |  |
| § Read coverage and NGx statistics have been calculated based on the estimated genome size of 315 Mb. |  |  |  |  |  |  |

During manual curation, we generated 8 breaks and 24 joins and we were able to close a total of 11. Finally, we filtered out 22 contigs from the assembly, with 21 corresponding to the endosymbiont, *Blochmannia*, and 1 corresponding to a mitochondrial contaminant. The Omni-C contact maps show that the assembly is highly contiguous (Figure 2C). We have deposited the resulting assembly on NCBI (See Table 2 and Data Availability for details).

### Endosymbiont genome assembly

The final *Blochmannia* genome (ypCanBloch1_iyCamVici1.0) is a single gapless contig with final size of 780,225 bp, which is close but not equal to the reference used as guide (ASM2358568v1; genome size = 783,921 bp). The base composition of the final assembly version is A=35.05%, C=13.94%, G=14.37%, T=36.64%. The bacterial genome presented here consists of 624 coding sequences, 39 transfer RNAs, 1 transfer-messenger RNA, 3 ribosomal RNAs, and 2 non-coding RNAs.

### Assembly comparisons

Genome metrics indicate that the bicolored carpenter ant assembly is highly contiguous (62 contigs, contig N50 of 15.9 Mb), with fewer contigs and a longer contig N50 than all currently available ant genomes (Figure 1C, Table S1). Although chromosome assignments were not determined for *C. vicinus*, 14 out of the 38 total scaffolds in the genome assembly approach sizes >15.1 Mb (MEAN +/- SD = 21.6 +/- 6.2 Mb), make up >99.6% of the genome assembly, and are comparable to the average chromosome sizes of genome assemblies from four representative ant species (MEAN +/- SD = 16.5 +/- 9.3 Mb, Figure 1D, Table S2).

### Phylogenetic analysis

Phylogenetic reconstruction placed our *C. vicinus* sample as a sister group to a clade including *C. vicinus* from Arizona and two individuals, also from Arizona, designated *C. sp*. (2-JDM) in Manthey *et al*. (2022). Given this result, placing these two *C. sp*. (2-JDM) in *C. vicinus* would restore monophyly for this species and yield a more inclusive, wide-ranging taxon. However, if closer morphological examination and population sampling reveal that these samples are not conspecific with *C. vicinus*, then the species may require further taxonomic rearrangement to resolve this species-level paraphyly.

## Discussion

The high-quality bicolored carpenter ant (*Camponotus vicinus*) genome assembly, presented here, will serve as a foundational reference for future evolutionary and population genomic studies in this and other related species. Our genome assembly is highly accurate, with coverage (46.8x) in range with other ant genome assemblies that include PacBio sequencing methods (coverage range: 45 - 245x, median coverage: 87x, Table S1) and BUSCO genome completeness (99.2%, compared with Arthropoda) slightly exceeds the median BUSCO values of other ant genome assemblies compared with the same BUSCO dataset (median BUSCO: 98.3%, BUSCO range: 68.0 - 99.6%, Table S1). In comparison with other ant genome assemblies, the bicolored carpenter ant assembly is the most contiguous (contig-level) assembly of all currently available ant genomes (Figure 1C, Table S1). Additionally, the 14 largest *C. vicinus* scaffolds compose 99.7% of the genome assembly, matching the predicted chromosome number of n=14 for *C. vicinu*s, based on the reported karyotypes of the related species *C. ligniperdus* and *C. japonicu*s (Hauschteck & Jungen 1983, Imai 1966), and are similar to the chromosome sizes of genome assemblies from four representative ant species (Figure 1D, Table S2). Taken together, these results indicate that our *C. vicinus* genome is a chromosome-level assembly.

In comparison to other *Camponotus* ant genome assemblies available for the Florida carpenter ant (*C. floridanus*, Shields *et al*. 2018) and the black carpenter ant (*C. pennsylvanicus*, Faulk 2023), our bicolored carpenter ant nuclear genome assembly is similar in size (302.7 Mb) to the black carpenter ant assemblies (306.4, haplotype 1; and 305.9, haplotype 2), which are respectively 6.6%, 7.9 %, and 7.7% larger than the Florida carpenter ant genome assembly (284.0 Mb). Additionally, the mitochondrial genome assembly of the bicolored carpenter ant (16,542 bp) is nearly identical in size to the black carpenter ant (16,536 bp). We also assembled the *Blochmannia* bacterial endosymbiont for *C. vicinus* (780,225 bp) whose size falls in range with assemblies of *Blochmannia floridanus* (705,557 bp, isolated from *C. floridanus*, Gil *et al*. 2003) and *Blochmannia pennsylvanicus* (791,499-791,654 bp, isolated from *C. pennsylvanicus*, Degnan *et al*. 2005, Faulk 2023). Lastly, phylogenetic analysis of the *C. vicinus* reference genome, in comparison to recently published whole genome sequences representing nine *Camponotus* species (Manthey *et al*. 2022, Shields *et al*. 2018, and Faulk 2023), revealed that *C. vicinus* (California, this study) is sister to the clade containing *C. vicinus* (Arizona) and *C. sp. 2-JDM* (Figure 1B). This analysis suggests that further investigation is needed to resolve the species assignment and resulting monophyly or paraphyly of these representative samples.

The reference genome of bicolored carpenter ant, *Camponotus vicinus*, will allow us to better understand the genetic basis of adaptations, track evolutionary changes, and assess genomic variation that may impact survival and speciation. Furthermore, the bicolored carpenter ant reference genome serves as a powerful tool for both evolutionary and conservation biologists to better understand the genetic makeup of the *C. vicinus* species complex, which can inform taxonomic studies of this group and contribute to efforts of the California Conservation Genomics Project (CCGP) (Shaffer et al. 2022). It fills an important phylogenetic gap in our genomic understanding of California biodiversity (Toffelmier et al. 2022). Future work comparing multiple genomes of *C. vicinus* across California will additionally help identify regions that are associated with species resilience and biodiversity, and aid in development of effective conservation and management strategies accordingly (Fiedler et al. 2022).

## Acknowledgements

PacBio Sequel II library prep and sequencing was carried out at the DNA Technologies and Expression Analysis Cores at the UC Davis Genome Center, supported by NIH Shared Instrumentation Grant 1S10OD010786-01. Deep sequencing of Omni-C libraries used the Novaseq S4 sequencing platforms at the Vincent J. Coates Genomics Sequencing Laboratory at UC Berkeley, supported by NIH S10 OD018174 Instrumentation Grant. We thank the staff at the UC Davis DNA Technologies and Expression Analysis Cores and the UC Santa Cruz Paleogenomics Laboratory for their diligence and dedication to generating high quality sequence data.

## Data Availability

The NCBI BioProject ID for the California Conservation Genomics Project is PRJNA720569. Data generated for this study are available under NCBI BioProject PRJNA808334. Raw sequencing data for sample RG_C_vicinus_PSW18456_S1, RG_C_vicinus_PSW18456_S2 (NCBI BioSample SAMN30501363,SAMN30501644) are deposited in the NCBI Short Read Archive (SRA) under SRX18986378 for PacBio HiFi sequencing data, and SRX18986379 for the Omni-C Illumina sequencing data. GenBank accession for the nuclear assembly is GCA_025532165.1 and JANXEZ010000038.1 for the mitochondrial assembly. Whole genome sequence accessions are under JANXEZ000000000.Assembly scripts and other data for the analyses presented can be found at the following GitHub repository: www.github.com/ccgproject/ccgp_assembly.

